# Integrated Sequencing & Array Comparative Genomic Hybridization in Familial Parkinson’s Disease

**DOI:** 10.1101/828566

**Authors:** Laurie A. Robak, Renqian Du, Bo Yuan, Shen Gu, Isabel Alfradique-Dunham, Vismaya Kondapalli, Evelyn Hinojosa, Amanda Stillwell, Emily Young, Chaofan Zhang, Xiaofei Song, Haowei Du, Tomasz Gambin, Shalini N. Jhangiani, Zeynep Coban Akdemir, Donna M. Muzny, Anusha Tejomurtula, Owen A. Ross, Chad Shaw, Joseph Jankovic, Weimin Bi, Jennifer E. Posey, James R. Lupski, Joshua M. Shulman

## Abstract

**Background:** Parkinson’s disease (PD) is a genetically heterogeneous condition; both single nucleotide variants (SNVs) and copy number variants (CNVs) are important genetic risk factors. We examined the utility of combining exome sequencing and genome-wide array-based comparative genomic hybridization (aCGH) for identification of PD genetic risk factors.

**Methods:** We performed exome sequencing on 110 subjects with PD and a positive family history; 99 subjects were also evaluated using genome-wide aCGH. We interrogated exome sequencing and array comparative genomic hybridization data for pathogenic SNVs and CNVs at Mendelian PD gene loci. SNVs were confirmed via Sanger sequencing. CNVs were confirmed with custom-designed high-density aCGH, droplet digital PCR, and breakpoint sequencing.

**Results:** Using exome sequencing, we discovered individuals with known pathogenic single nucleotide variants in *GBA* (p.E365K, p.T408M, p.N409S, p.L483P) and *LRRK2* (p.R1441G and p.G2019S). Two subjects were each double heterozygotes for variants in *GBA* and *LRRK2*. Based on aCGH, we additionally discovered cases with an *SNCA* duplication and heterozygous intragenic *GBA* deletion. Five additional subjects harbored both SNVs (p.N52fs, p.T240M, p.P437L, p.W453*) and likely disrupting CNVs at the *PARK2* locus, consistent with compound heterozygosity. In nearly all cases, breakpoint sequencing revealed microhomology, a mutational signature consistent with CNV formation due to DNA replication errors.

**Conclusions:** Integrated exome sequencing and aCGH yielded a genetic diagnosis in 19.3% of our familial PD cohort. Our analyses highlight potential mechanisms for *SNCA* and *PARK2* CNV formation, uncover multilocus pathogenic variation, and identify novel SNVs and CNVs for further investigation as potential PD risk alleles.

## Introduction

Parkinson’s disease (PD) is the second most common adult-onset neurodegenerative disorder with substantial evidence for heritability.^1–3^ Up to 20% of patients with PD report a positive family history^4–6^ and genetic risk factors are more common in these families.^7^ More than 40 different loci that increase PD susceptibility have been identified in familial and sporadic PD^8–12^. Although PD is diagnosed based on clinical criteria,^13^ identification of specific genetic risk factors can reveal prognostic information, such as risk of cognitive impairment and/or rate of progression,^14, 15^ and may soon highlight eligibility for personalized therapies.^16, 17^ In addition, discovery of risk variants may inform genetic counseling of unaffected family members. Indeed, surveys of patients and caregivers reveal a high level of interest in genetic testing for PD.^18, 19^

Exome sequencing (ES) has accelerated the discovery of PD genetic risk factors^20–22^ and is ideally suited to identify single nucleotide variants (SNVs) in genetically heterogeneous diseases. In one study of adult patients referred for diverse clinical indications, ES had a diagnostic yield of 10% in individuals over 30 years of age.^23^ In a recent study of 80 early-onset sporadic PD cases, ES yielded an overall diagnostic rate of 11%, with *GBA* alleles accounting for 5%.^24^ Nevertheless, clinical genetic testing is not routinely performed for PD, and ES remains poorly studied as a potential genetic diagnostic tool.

Although most identified PD risk alleles are SNVs, chromosomal structural rearrangements, or copy number variants (CNVs), also play an important role.^25^ Despite notable recent advances,^26^ ES remains insensitive for detection of small CNVs (< 50Kb).^27^ Several complementary approaches, including multiplex ligation-dependent probe amplification.^28^ bacterial artificial chromosome arrays,^29^ and single nucleotide polymorphism arrays^30, 31^ have shown mixed success for identification of CNVs in PD cohorts. In contrast, genome-wide array-based comparative genomic hybridization (aCGH), is a highly-validated, sensitive clinical screening tool for CNV detection, offering exon-by-exon coverage for a multitude of disease-associated genes.^32–35, 34, 35^ While not yet adopted in most diagnostic laboratories, droplet digital Polymerase Chain Reaction (ddPCR) is also emerging as a rapid and cost-efficient, targeted approach for the assessment of small CNVs at specific loci.^36^ Compared to standard quantitative PCR, digital PCR offers enhanced copy number and gene dosage sensitivity, precision, and reliability due to sample partitioning.^37^

We explored the genetic molecular diagnostic rate for integrated ES and aCGH in a familial PD cohort. In addition, we evaluate ddPCR for confirmation of pathogenic CNVs, and using breakpoint sequencing, we elucidate potential mechanisms for CNV formation.

## Methods

### Subjects

We studied 110 PD cases evaluated in the Baylor College of Medicine (BCM) PD Center and Movement Disorders Clinic in Houston, TX with a family history of PD (45% of cases reported an affected first degree relative; 55% reported a second or third degree relative). All diagnoses were made by movement disorder specialists. Subjects in our cohort were unrelated, except for 2 brothers (subjects 21 and 22) known to have *PARK2*-related PD.^12^ However, neither ES nor aCGH were previously performed in this sibling pair. Parental and other family member samples were not available for any subjects. All subjects provided informed consent and the study was approved by the BCM Institutional Review Board. As a positive control for aCGH, we also included a sample from a known subject with an *SNCA* triplication.^38–41^ We also interrogated a Baylor Genetics diagnostic laboratory sample including 12,922 clinical referral samples for aCGH from peripheral blood.^42^ This analysis of aggregate clinical genomic data was also approved by the BCM Institutional Review Board. Subject numbers throughout the text are consistent with clinical and demographic details provided in Supplementary Table 1.

### Gene Set Definition and Variant Criteria

We focused our analyses on genes and variants established to cause familial PD, including the autosomal dominant loci, *SNCA* (MIM#168601),^43^ *GBA* (MIM#168600),^44, 45^ *LRRK2* (MIM#607060),^46–48^ *GCH1* (MIM#600225),^20^ *DNAJC13* (MIM#616361),^49, 50^ and *VPS35* (MIM#614203),^51, 52^ and the autosomal recessive loci, *PARK2* (*PRKN,* MIM#600116),^53^ *PINK1* (MIM#605909)^54^ and *PARK7* (*DJ1*, MIM#606324),^55^ based on the available literature in April 2015 when this study was initiated. Deletions of 22q11.2 have recently been recognized as incompletely-penetrant PD risk factors^56, 57^ and were also considered for CNV analyses. From the ES data, all candidate variants were reviewed by Clinical Geneticists (LR, JP, JL) and Movement Disorder Neurologists (JS, JJ) to establish a consensus on pathogenic alleles, integrating data from multiple available resources (PubMed, ClinVar^58^, Human Genome Mutation Database^59^). All pathogenic alleles included in this study are well-established, non-synonymous coding variants with moderate to high penetrance (OR>2) meeting stringent evidence for replication across studies or within the same study. All other variants discovered in these genes but not previously reported in PD were considered variants of unknown significance (VUSs).

### SNV and CNV Detection

Genomic DNA was extracted from peripheral blood samples obtained from each participant. ES for the first 50 subjects was obtained from a prior study^21^, and ES for the remaining 60 subjects was performed at the BCM Human Genome Sequencing Center (HGSC) using the Illumina HiSeq 2000.^60^ Samples achieved an average of 95% of targeted exome bases covered to a depth of 20X or greater. All pathogenic SNVs detected via ES were confirmed via Sanger sequencing. Genome-wide array CGH was performed on 99 out of 110 subjects for which sufficient DNA remained after ES, using Baylor Genetics V10 2×400K high-density clinical-grade oligonucleotide microarrays, which were previously used for clinical diagnostic purposes. We defined potential CNVs as those regions with three or more consecutive probes with consistent direction of effect. To confirm these potential CNVs, we employed a custom 8×60K high-density array through Agilent (CA, USA). For the Baylor Genetics clinical cohort lookup, data was available from 12,922 subjects profiled for CNVs using Baylor arrays (v9 or v10).^42^ CNVs were filtered according to the following criteria: 3+ consecutive probes with consistent direction of effect, log_2_ ratios for loss <=0.5, gain>0.35. Filtered CNVs were manually assessed to exclude artifacts.^61^ For comparison of CNV frequencies, the 2-tailed chi square test was employed. See also Supplemental Methods for further details, including ddPCR protocol.

## Results

We pursued genetic diagnostic evaluation of 110 total subjects (including 109 unrelated probands) with familial PD. The mean age at onset was 50 years (SD 15); 51% were male. The ethnic composition of the cohort was 72% Caucasian, 17% Hispanic, 6% East Asian, South Asian or Middle Eastern, and 6% undefined (not reported).

### Single Nucleotide Variants

We first examined subject ES data for pathogenic SNVs in established PD genes (see Methods). Among the dominant PD loci, 15 individuals had variants in *LRRK2* (c.6055G>A:p.G2019S and c.4321C>G:p.R1441G) and *GBA* (c.1093G>A:p.E365K, c.1223C>T:p.T408M, c.1448T>C:p.L483P, and c.1226A>G:p.N409S) (Table 1). Two subjects each harbored heterozygous SNVs in both *GBA* and *LRRK2*; i.e. double heterozygotes. One subject had a combination of *LRRK2* p.G2019S and *GBA* p.E365K (Subject 2), while the other had *LRRK2* p.G2019S and *GBA* p.L483P (Subject 13). Both subjects had onset of PD symptoms in their 40s. On initial exam, Subject 2 had tremor at rest, rigidity, bradykinesia and dystonic posturing in both hands. She reported a history of PD in her father and paternal grandfather (Supplementary Figure 1A; Supplementary Table 1). There was no history of cognitive impairment or dementia. Subject 13 presented with resting tremor, rigidity and bradykinesia. She reported a family history of PD in her paternal uncle (Supplementary Figure 1B, Supplementary Table 1). Ten years after PD diagnosis, she developed visual hallucinations and delusions. The subjects were of European and Hispanic ancestry, respectively; neither reported Ashkenazi Jewish heritage. ES also revealed seven individuals with pathogenic variants in loci usually associated with autosomal recessive PD, including *PARK2* (6 individuals) and *DJ-1/PARK1* (1 individual). However, all subjects were heterozygous SNV carriers, and therefore isolated ES was non-diagnostic (Table 1). Therefore, based on ES alone, we identified a pathogenic variant accounting for PD in 13.8% (n=15 of 109 probands) of our familial PD cohort. All implicated variants were confirmed via Sanger sequencing. Besides the pathogenic variants noted above, ES also identified heterozygous variants of unknown significance (VUS) in many PD risk genes (Supplementary Table 2), including *LRKK2* (n=12), *GCH1* (n=1), *VPS35* (n=1), *DNAJC13* (n=18), *DJ-1/PARK7* (n=2)*, PARK2* (n=9), and *PINK1* (n=1). For recessive PD genes, all subjects harbored a single heterozygous variant.

**Table 1:** SNVs associated with increased PD risk detected via exome sequencing. All indicated SNVs were heterozygous, except *PARK2* c.719C>T:p.T240M, which was hemizygous, as the variant is in *trans* to a deletion allele. Pathogenic variants were considered diagnostic (Y) if discovered in an autosomal dominant gene, or in the case of autosomal recessive genes, if in combination with a CNV (asterisk, see also Figure 3). Non-diagnostic (N), heterozygous SNVs were also discovered in *PARK2* (p.R275W) and *DJ-1* (p.A104T). In 2 subjects, SNVs in both *LRRK2* and *GBA* were identified (double heterozygotes): ^A^*LRRK2* p.G2019S and *GBA* p.E365K; ^B^*LRRK2* p.G2019S and *GBA* p.L483P. ^C^Interpretations of c.1310C>T:p.P437L are conflicting ^94, 95^.

| Gene | Variant | Diagnostic | Subjects (n) |
| --- | --- | --- | --- |
| <u>Dominant</u> |  |  |  |
| *LRRK2* | c.6055G>A:p.G2019S | Y | 3<sup>A,B</sup> |
| *LRRK2* | c.4321C>G:p.R1441G | Y | 1 |
| *GBA* | c.1093G>A:p.E365K | Y | 5<sup>A</sup> |
| *GBA* | c.1223C>T:p.T408M | Y | 5 |
| *GBA* | c.1448T>C:p.L483P | Y | 1<sup>B</sup> |
| *GBA* | c.1226A>G:p.N409S | Y | 2 |
| <u>Recessive</u> |  |  |  |
| *PARK2* | c.155delA:p.N52fs | Y* | 1 |
| *PARK2* | c.1310C>T:p.P437L<sup>C</sup> | Y* | 1 |
| *PARK2* | c.1358G>A:p.W453* | Y* | 1 |
| *PARK2* | c.719C>T:p.T240M | Y* | 2 |
| *PARK2* | c.823C>T:p.R275W | N | 1 |
| *DJ-1* | c.310G>A:p.A104T | N | 1 |

### Copy Number Variants

We next interrogated aCGH data for pathogenic CNVs among PD genes. Our analyses included 99 out of 110 total subjects evaluated by ES. No CNVs were observed in *VPS35, LRRK2, DNAJC13, GCH1, DJ1,* or *PINK1*, nor did we identify any candidate deletions at the 22q11.2 locus. However, we discovered CNVs in *SNCA* (n=1), and *GBA* (n=1), and *PARK2* (n=5). All reported CNVs were confirmed by custom high-density arrays and breakpoint sequencing. We first discuss CNVs in *SNCA* and *GBA* since these affect dominant PD genes and were considered diagnostic based on aCGH alone. All heterozygous CNVs in the recessive PD gene, *PARK2*, were considered further in combination with ES results (see next section). Overall, isolated aCGH identified a genetic risk factor for PD that was diagnostic in 2.0% of our cohort (n=2 of 99 probands). Based on aCGH, we have also detected numerous large CNVs (> 1 Mb) within our cohort that impact other genomic loci; these variants remain of uncertain clinical significance but are documented for completeness (Supplementary Table 3).

In Subject 3, we detected a 248 kb duplication encompassing *SNCA*, as well as the adjacent *MMRN1,* which has no known association with human disease. This CNV was confirmed by high-density aCGH and breakpoint analysis (Figure 1A). On initial exam, this subject exhibited rigidity, tremor, gait impairment, along with hyperreflexia and clonus. The subject was of Hispanic and Native American ancestry; the subject’s father had PD with dementia. Besides providing independent confirmation, breakpoint sequencing can provide clues to mechanisms of CNV formation. In the case of Subject 3, we identified a 1-bp microhomology domain, which is a short sequence that is identical to another region in the genome reduced from two copies to one during the template switch accompanying replicative repair.^62, 63^ Microhomology has been reported as a signature for a class of DNA replication errors that can generate CNVs (see Discussion).^64, 65^ As a positive control for our aCGH analysis, we also included a known *SNCA* triplication sample from the index family in which *SNCA* locus multiplication was first discovered as a cause for PD.^38–41^ Breakpoint sequencing revealed that this copy number alteration is a 1.7 Mb complex genomic rearrangement (Figure 1B), consisting of a duplication-inverted triplication-duplication (DUP-TRP/INV-DUP). This finding confirms and extends prior investigation of this particular structural variant^66^ and is also consistent with a likely replication-based mechanism for CNV formation.^67^

**Figure 1:**
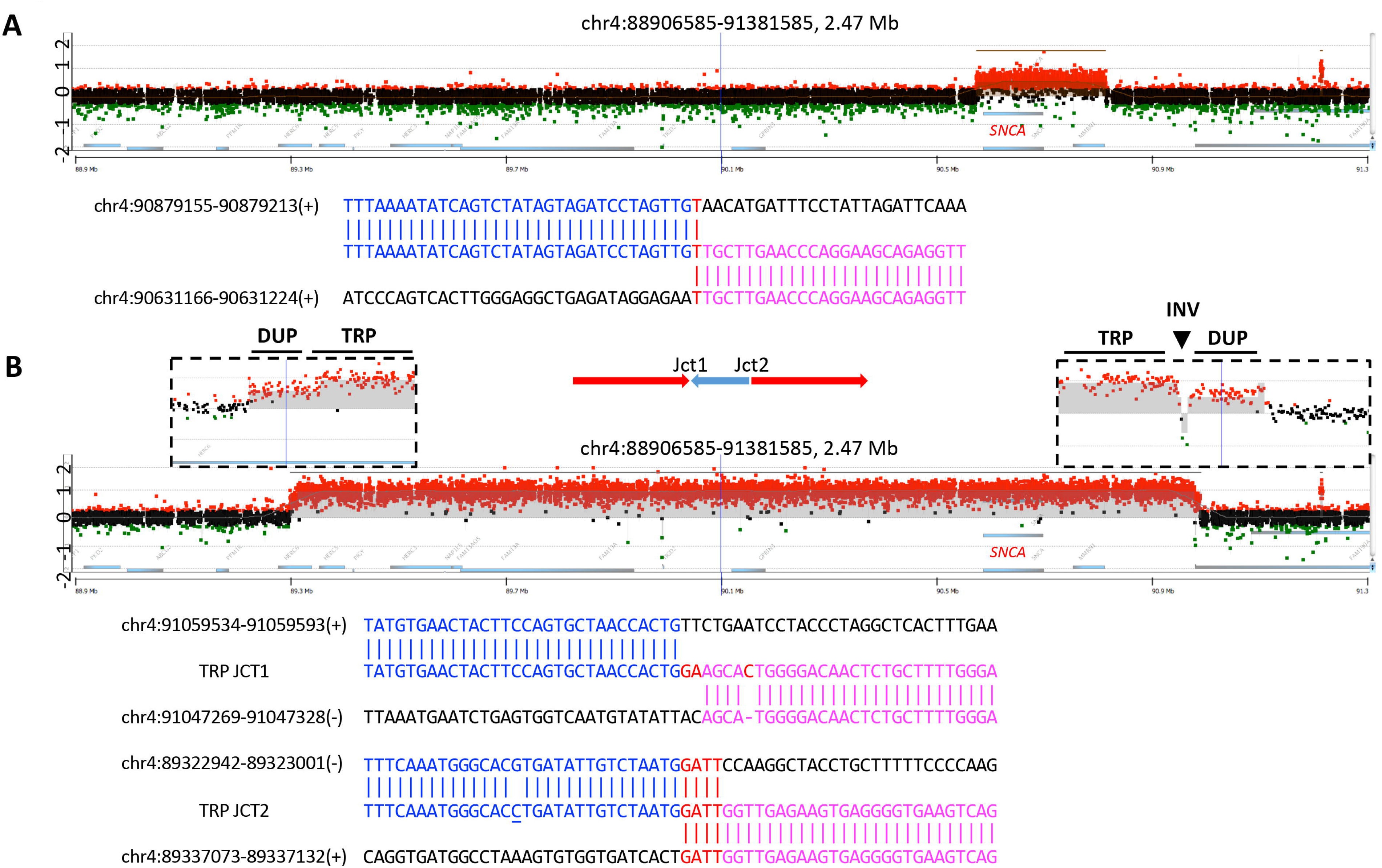
aCGH plots and breakpoint junction sequences of two CNVs involving *SNCA*. (A) In Subject 3, a 248-kb duplication was identified. In this case, the whole *SNCA* gene was duplicated. The junction sequence (Bottom) is aligned with upstream and downstream reference sequences, with the blue and pink colors indicating their different origins, and the red indicating inserted nucleotides and microhomology. (B) A 1.7-Mb DUP-TRP/INV-DUP rearrangement was identified in the index patient with a known *SNCA* multiplication^39, 40, 82^ The x-axis indicates the chromosomal regions surrounding *SNCA*. The y-axis indicates the subject versus control log_2_ ratio of the aCGH results, with duplications at 0.58, triplications at 1, and heterozygous deletions at −1 based on theoretical calculations. Red dots in the graph represent probes with log_2_ ratio >0.25, black dots with log_2_ ratio from 0.25 to −0.25, and green dots with log_2_ ratio <-0.25. The normal-duplication-triplication transition regions are magnified in boxes above the plot. The entire *SNCA* gene is triplicated. In addition, a SNP (rs12651181, underlined) was detected close to JCT2.

We also discovered a heterozygous intragenic deletion in *GBA* in one subject (Figure 2), who presented at age 28 with tremor, bradykinesia and rigidity. She had an excellent response to levodopa in her early 30s, and subsequently developed dyskinesia. The subject was of European descent, and both of her maternal grandparents were also diagnosed with PD (Supplementary Figure 1C). Since variant confirmation at the *GBA* locus can be complicated by an adjacent pseudogene with significant homology,^68^ we confirmed the 4.7 kb deletion of exons 2-8 using long-range PCR (Figure 2C). Breakpoint sequencing also confirmed heterozygosity and further revealed a 5-bp microhomology domain consistent with CNV formation due to non-homologous recombination or replication errors (Figure 2B).

**Figure 2:**
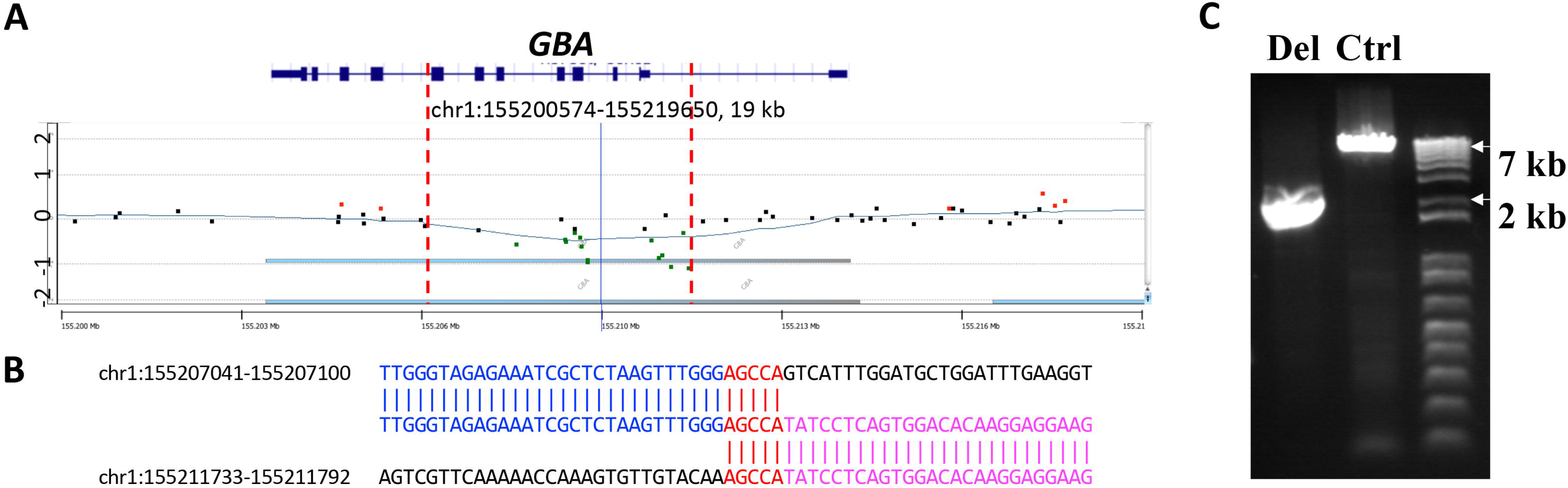
aCGH plot and breakpoint junction sequence of *GBA* deletion. (A) aCGH plot and (B) junction sequence of a 4.7-kb deletion identified involving *GBA* in patient 1. The deletion (shadowed) encompasses 7 exons of *GBA* (from exon 2 to exon 8). (C) By agarose gel electrophoresis, the amplification of the deleted region (Del) in Subject 1 showed a ∼5kb discrepancy compared to a control subject (Ctl), consistent with aCGH findings. PCR showed preferential amplification of the shorter fragment in the Del lane.

### Integrated Analysis of SNVs and CNVs

As highlighted above, isolated ES and aCGH each discovered a number of subjects with heterozygous SNVs or CNVs affecting recessive genes. In order to determine if these changes might be diagnostic, we examined the results together for potential biallelic variation at the locus due to both a SNV allele and a CNV allele. Indeed, three subjects in our cohort were newly identified as potential compound heterozygous carriers of both a pathogenic CNV and SNV in *PARK2* (Figure 3). Subject 6 had a 364 kb duplication of exons 4-6 and a frameshift deletion c.155delA:p.N52fs. Subject 20 had a 222 kb deletion of exons 8-9 and a stopgain c.1358G>A:p.W453*. Subject 11 harbored a pathogenic *PARK2* SNV (c.1310C>T:p.P437L) and a complex locus rearrangement, including a copy number neutral region flanked by 404 kb and 199 kb duplications affecting exons 2-3 and exons 5-6, respectively (Figure 3B). Our cohort also included 2 brothers with known *PARK2* PD;^12^ however, neither ES nor aCGH were previously performed. Our analysis confirmed compound heterozygosity for the known pathogenic SNV in exon 6 (c.719C>T:p.T240M, apparently homozygous on ES) and a CNV (178-kb deletion of exons 5-6) (Figure 3C). Interestingly, ES also discovered an additional VUS (c.2T>C:p.M1T). Based on available clinical information (Supplementary Table 1), all subjects with *PARK2* variants had young-onset PD (age range 15-36).

**Figure 3:**
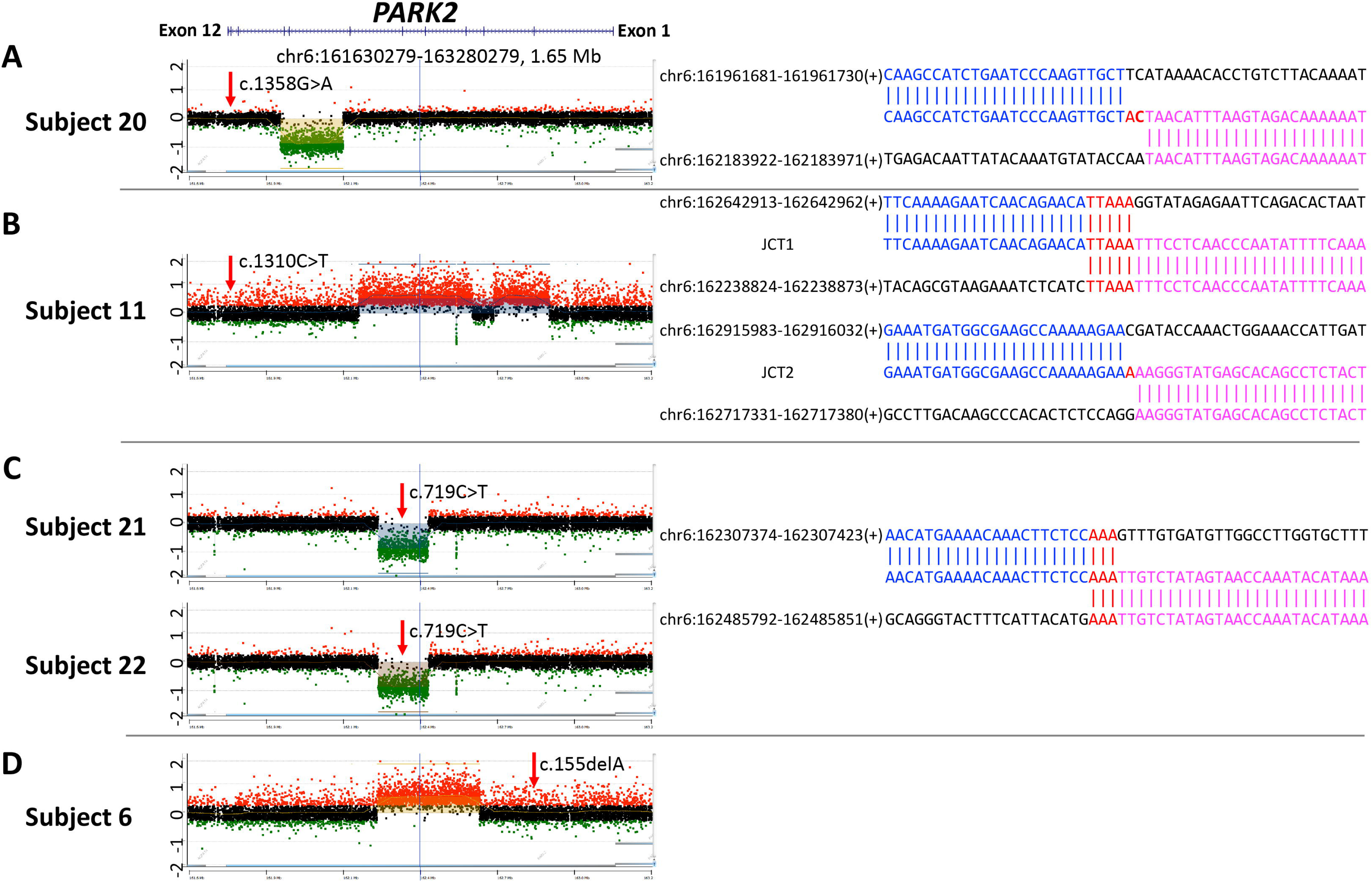
aCGH plots and breakpoint junction sequences of *PARK2* CNVs. aCGH plots (left panel) and breakpoint junction sequences (right panel) of CNVs identified involving *PARK2* gene in the cohort. At the top, a schematic gene structure demonstrates the 12 exons of *PARK2*. (A) In subject 20, a 222-kb deletion covering exons 8 and 9 was accompanied by a known pathogenic nonsense mutation c.1358G>A:p.W453* (gnomAD frequency = 0) in exon 12. (B) In subject 11, in addition to a missense variant c.1310C>T:p.P437L (exon 12)) a duplication-normal-duplication (DUP-NML-DUP) was identified. (C) Siblings 21 and 22 share a pathogenic missense variant c.719C>T:p.T240M (in exon 6) and a 178 kb deletion (disrupting exons 5 and 6). (D) In subject 6, a 364-kb duplication encompassed exons 4 to 6. A known pathogenic frameshift mutation c.155delA:p.N52fs was identified in exon 2 (gnomAD frequency = 2.5×10^-4^). Breakpoint sequencing was not successful in this sample.

All CNVs were again confirmed using custom, high-density arrays as well as breakpoint sequencing. In the case of subject 6, we were unable to successfully amplify breakpoint junctions despite multiple attempts, suggesting a more complex genomic rearrangement or raising the possibility that this duplication is located elsewhere in the genome. For all other *PARK2* CNVs, junction structure is detailed in Figure 3, including implicated mechanisms of CNV formation. Overall, integrated ES and CNV identified a genetic cause for PD in 4 additional probands, increasing the overall genetic diagnostic yield to 19.3% (n=21).

### Digital Droplet PCR (ddPCR)

Given the observed high frequency of *PARK2* CNVs in this familial PD cohort (∼5%), and the high cost of clinical-grade aCGH, we examined the feasibility of exon-by-exon ddPCR assay to detect *PARK2* CNVs as a proof of principle. ddPCR is an emerging cost-effective method for sensitive, and reliable assessment of specific CNVs^36, 37^. Indeed, all *PARK2* CNVs described (subjects 6, 11, 20, 21 and 22) were robustly detected using ddPCR (Figure 4A). An additional 94 cases from our cohort with available DNA were interrogated for intragenic *PARK2* CNVs, and no additional CNVs were detected (Figure 4A and Supplementary Figure 2). We also applied ddPCR to screen for potential CNVs at the *GBA* locus. The 12-exon *GBA* shares high homology with a nearby 13-exon pseudogene, *GBAP1*. As shown in Figure 4C, using ddPCR, we successfully amplified all 12 exons of *GBA*. Six exons (1, 2, 4, 7, 9, and 10) identified unique amplicons with a positive droplet ratio of 1, while the other six exons (3, 5, 6, 8, 11, and 12) demonstrated shared amplicons with the pseudogene, resulting in a droplet ratio of 2. Importantly, the deletion of *GBA* exons 2-8 in Subject 1 was successfully detected and confirmed by ddPCR (Figure 4C), and the assay was negative for *GBA* CNVs in 86 other samples tested (Figures 4C and Supplementary Figure 3). Of note, a single exon deletion (exon 6) was suggested by ddPCR in subject 48. Further investigation revealed an intronic SNV located at the 3’ end of one ddPCR primer affecting amplification efficacy, confirmed by Sanger sequencing (Supplementary Figure 4A). We re-designed the affected primer (Supplementary Methods) and demonstrated full amplification of exon 6 (Supplementary Figure 4B). Overall, our results suggest that ddPCR may be a sensitive and specific diagnostic tool for CNV detection in PD subjects, including at loci such as *GBA* complicated by genomic regions with high sequence homology.

**Figure 4:**
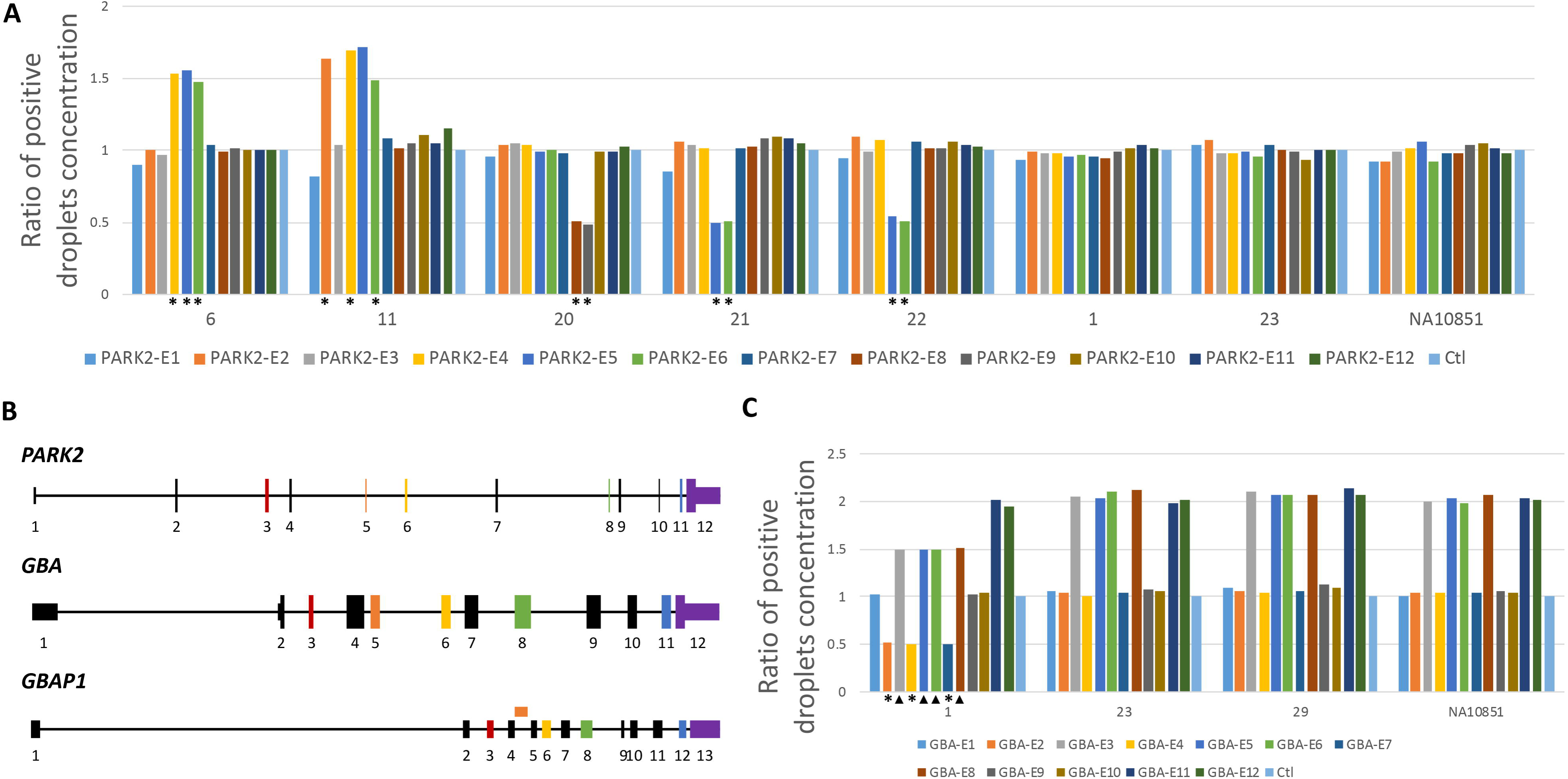
*PARK2* and *GBA* ddPCR results of representative subjects. (A) Positive droplet concentrations in 8 subjects are shown. Primer pairs for the 12 exons of *PARK2* and two control genes, *RPPH1* and *TERT,* were used to obtain positive droplet concentrations from PCR in each individual (Supplementary Methods and Supplementary Figure 2). The y-axis shows exon-by-exon results in 13 columns with different colors, showing comparable results to the average value of *RPPH1* and *TERT*.A y-axis value of 0.5 indicates a deletion, 1 copy neutral (no deletion, no duplication), and 1.5 a duplication. In subject 6, a duplication involving exons 4 to 6 was identified as shown by aCGH,; in subject 11, exons 2, 4, 5, and 6 demonstrated copy number gains; in subject 20, there is a copy number loss involving exons 8 and 9; similarly, in subjects 21 and 22 a copy number loss of exons 5 and 6 is detected. In subjects 1, 23, and HapMap NA10851, no amplicons showed altered copy number. See also Supplementary Figure 2. Copy number variants are denoted with asterisks (*). (B) *GBA* and its nearby pseudogene, *GBAP1*, share a high degree of sequence homology, with ddPCR primer pairs for 6 of the 12 exons of *GBA* producing amplicons concurrently from *GBA* and *GBAP1*: *GBA* exons 3, 5, 6, 8, 11, and 12 are color coded to demonstrate their homologous regions within *GBAP1,* which result in a doubling of the apparent copy number identified by ddPCR: 4 instead of 2 copies (ratio = 2), indicate copy number neutrality for these exons. *GBA* exon 5 is homologous with an intragenic region between exons 4 and 5 of *GBAP1*. (C) ddPCR was used to detect potential exonic CNVs in *GBA*. Here, we demonstrate a deletion identified in subject 1, compared to HapMap subject NA10581 and other two subjects, ratios of exons 2 to 8 were each reduced by 0.5-fold, consistent with a deletion involving these exons. Deleted exons are denoted with an asterisk (*); deleted exons with a droplet ratio of 1.5 due to *GBAP1* amplification are denoted with an arrowhead.

### CNV Burden in Clinical Cohorts

Compared to SNVs, limited reference data is available on the population frequency of CNVs, especially in neurologically healthy adult samples, hampering the interpretation of CNV frequencies detected in our cohort. We therefore leveraged data from the Baylor Genetics diagnostic laboratory, including 12,922 aCGH clinical referral samples. This large cohort is skewed for pediatric cases (mean age=7.4 years, SD = 9.7 years, range 0-79 years), reflecting the more common use of aCGH in this population. Although the cohort includes a substantial proportion of individuals with developmental delay, autism and dysmorphic features, there were no recorded submissions for PD. Based on stringent criteria (see Methods), at most PD loci, CNVs were either absent (*DNAJC13, LRRK2, PINK1,* and *SNCA*) or very rare at *VPS35* (n=2)*, GCH1* (n=1), and *DJ-1* (n=6, all subjects had 1p36 deletion syndrome). By contrast, CNVs were more common at 22q11.2 (n=90, all losses affecting the critical region) and *PARK2* (n=95). Notably, the frequency of *PARK2* CNVs in our PD cohort (5.1%) represents a significant enrichment (*p*=8.5×10^-7^), when compared to that of the Baylor Genetics clinical reference sample (0.74%). Due to the pseudogene, *GBAP1*, and suboptimal probe coverage, the array does not reliably capture *GBA* CNVs.

## Discussion

Establishing a specific genetic diagnosis can provide information about PD risk and progression relevant to patients and their families, and may soon influence treatment decisions.^17, 69^ We have evaluated combined ES and aCGH for genetic diagnosis in 110 subjects with familial PD (both assays were available for 99 subjects). In our cohort, ES and aCGH independently identified a genetic cause for PD in 13.8% and 2.0%, respectively. Our diagnostic yield for ES was slightly higher than that recently reported for an early-onset PD cohort (11.25%),^24^ and was also greater than the 10.7% diagnostic rate in an unselected adult series referred for clinical diagnostic ES.^23^ Given incipient treatment trials for *GBA-PD* and the potential importance of identifying eligible subjects in the future^69^, we included lower-risk pathogenic alleles (OR∼2.4)^70^, p.E365K and p.T408M, along with higher-penetrance variants (e.g. p.L483P, OR>5)^44^. Importantly, integrated ES and aCGH identified 5 additional subjects (4 unrelated probands)—including a subject with a *GBA* deletion—yielding an overall combined diagnostic rate of 19.3%. Our analyses also uncovered numerous VUS, including SNVs within Mendelian PD genes (Supplementary Table 2) as well as large CNVs affecting other loci (Supplementary Table 4). While additional evidence will be required to confirm or refute pathogenicity, our genetic diagnostic rate would nearly double if the variants in PD genes are considered *bona fide* risk factors. Overall, our findings suggest that complementary assessments for SNV and CNVs will be essential for routine, high-confidence genetic diagnosis in familial PD.

Most genetic diagnostic studies in PD cohorts to date have ignored the potential contribution of CNVs. Similarly, except in several notable targeted CNV studies, ^29, 30, 56, 71^ research-based PD gene discovery has almost exclusively focused on SNVs, using ES or genotyping arrays.^22, 72^ In our familial PD cohort, pathogenic alleles at both autosomal dominant (*SNCA*, *GBA*) and recessive (*PARK2*) loci would have been missed if not for the inclusion of aCGH for detection of CNVs. For example, out of 7 pathogenic alleles discovered using ES at recessive loci (*PARK2* and *DJ-1*), nearly all cases (except subjects 21 and 22) would have been non-diagnostic leading to misclassification as heterozygous carriers without aCGH. However, integrated SNV-CNV analyses successfully resolved 5 individuals with likely *PARK2* PD. Our findings suggest caution for interpretation of studies attributing PD risk to either *PARK2* CNV or SNV heterozygous carrier states in isolation, consistent with prior studies.^73, 74^ The importance of integrated SNV-CNV analysis likely extends to other autosomal recessive PD loci, including *DJ-1* and *PINK1.*^55, 75^ Since our CNV and SNV data are unphased, and parental genotypes are not available, we cannot definitively exclude the possibility that certain CNVs and SNVs at *PARK2* were in *cis-* rather than *trans-* configuration. Nevertheless, our data suggest that structural variants may co-occur with SNVs more commonly than previously recognized, making consideration of both allele types important for comprehensive genetic diagnosis in PD.^76^

ES has significantly accelerated the scope of gene discovery in PD and other neurologic disorders,^22, 24^ but remains insensitive to allele classes such as trinucleotide repeat expansions and CNVs. While bioinformatic tools can be employed to screen for CNVs in data from ES, available algorithms have high false positive rates compared to aCGH^77^ and as many as 30% of clinically relevant CNVs are missed by ES.^27^ To our knowledge, genome-wide aCGH with exon-by-exon coverage has not been previously applied in PD. Limitations of aCGH include significant cost, and the possibility of missing small deletions/duplications. Alternative methods, such as ddPCR, may offer a cost effective alternative for screening specific genes,^36^ including for small CNVs. In our study, ddPCR showed high sensitivity and specificity for detection of CNVs at both *PARK2* and *GBA*. In the latter case, ddPCR was able to robustly differentiate copy number changes affecting exons unique to *GBA* avoiding potential confounding by the adjacent pseudogene, *GBAP1*.

Despite evidence of an important role in disease risk, the mechanism(s) responsible for generating CNVs relevant to PD have been poorly studied. Broadly, CNVs may form through mechanisms associated with DNA recombination, DNA replication, and/or DNA repair.^62^ Mechanisms such as nonallelic homologous recombination^78–80^ can result in recurrent rearrangements. In contrast, non-homologous end joining,^64, 81^ fork stalling and template switching (FoSTeS) and microhomology-mediated break-induced replication (MMBIR),^64^ lead to non-recurrent CNVs. In our study, for all CNVs for which breakpoint junctions were obtained, the mechanism was consistent with FoSTeS/MMBIR. This finding has important implications for screening assays since methods sensitive for heterogeneous, exon-by-exon changes must be employed to detect non-recurrent CNVs. The FoSTeS/MMBIR mechanism can also trigger multiple iterative template switches in a single event, leading to the generation of more complex genomic rearrangements. Breakpoint sequencing of an *SNCA* CNV first observed in the Spellman-Muenter/Iowa kindred,^40, 82^ confirmed the DUP-TRP-DUP structure^66^ and further revealed an internal inversion (DUP-TRP/INV-DUP) and also revealed microhomology. This rearrangement must have arisen during mitosis via FoSTeS/MMBIR^62, 64, 83, 84^ and therefore likely represents a *de novo* triplication, in contrast to the meiotic *PMP22* triplications observed in Charcot Marie Tooth (MIM#118220), which derives from a duplication in the previous generation.^85^ Our results therefore demonstrate the essential role of breakpoint junction sequencing in definitively resolving CNV structure and responsible mechanisms.

To our knowledge, our study includes the first report of an intragenic *GBA* deletion allele in a subject with PD. Similar deletions in *GBA* have been rarely described in autosomal recessive Gaucher disease (MIM#230800).^86–88^ Our discovery of a *GBA* deletion allele, expected to cause glucocerobrosidase haploinsufficiency, adds to other emerging evidence supporting a loss-of-function mechanism in *GBA-*associated PD.^89, 90^ It will be informative to screen for additional *GBA* CNVs in additional case/control cohorts—perhaps using ddPCR—to determine how commonly these alleles are associated with PD risk and estimate their effect size and penetrance. We also identified 2 subjects in our cohort doubly heterozygous for SNVs in both *GBA* and *LRRK2*, and similar PD cases have been previously reported.^91^ As comprehensive, genome-wide diagnostic approaches, including ES and aCGH, become widely utilized in PD, it is likely that additional cases compatible with oligogenic inheritance models will be recognized.^92, 93^ Future studies will be required to define whether and how such alleles may interact to modify PD risk and/or clinical manifestations. Finally, while our study focused on pathogenic alleles in established, Mendelian loci, future assessment of a more complete spectrum of genetic variation through integrated SNV-CNV analysis is also likely to enhance power for novel PD gene discovery.

## Authors’ Roles

1. Research project: A. Conception: LAR, JRL, JMS B. Organization: LR, JRL, JEP, JMS C. Execution; LAR, RD, BY, SG, IAD, VK, EH, AS, EY, CZ, XS, HD, TG, SNJ, ZCA, DMM, AT, OAR, CS, JJ, WB, JRL, JMS
2. Statistical Analysis: A. Design: LR, WB, JEP B. Execution: LR, WB, CS, JEP C. Review and Critique:; LAR, RD, BY, SG, IAD, VK, EH, AS, EY, CZ, XS, HD, TG, SNJ, ZCA, DMM, AT, OAR, CS, JJ, WB, JRL, JEP, JMS
3. Manuscript: A. Writing of the first draft: LAR, RD, VK, JEP, JRL, JMS. Review and Critique: LAR, RD, BY, SG, IAD, VK, EH, AS, EY, CZ, XS, HD, TG, SNJ, ZCA, DMM, AT, OAR, CS, JJ, WB, JRL, JEP, JMS

## Acknowledgements

We thank all of the families who participated in this study. We also thank Drs. Dennis W. Dickson and Zbigniew Wszolek for providing the *SNCA* positive control sample. We acknowledge Mitchell A. Rao and Ido Machol for data retrieval from the clinical database.

## Supplementary Material

Supplementary material includes methods, 3 tables, and 4 figures.

