## Supplemental File for "Integrated Sequencing & Array Comparative Genomic Hybridization in Familial Parkinson’s Disease"

**Supplementary Methods.** The following primer sequences were used for breakpoint sequencing and ddPCR. *Adopted from Gu *et al*[1]; **Re-designed primer for exon 6 of *GBA*.

**Breakpoint Sequencing**

| Primer | Sequence (5' to 3') |
|---|---|
| SNCA-TRP-R | TCCTGTTCGGTTTTAGTCTGATGAAGCA |
| SNCA-DUP-R | GCCTCCACTGCACTTGTTATGAGCCA |
| SNCA-TRP-F | GCCAGCATATGACAAAACCCT |
| SNCA-DUP-F | ATGGTGAGCTCTATGAGGGC |
| 3-SNCA-F | TGCTTATTTCCTGTCCCGGT |
| 3-SNCA-R | TCAACCCAATCAACTGCAGC |
| 1-GBA-F | GTAGGAATCCTGGAGTTGGGTGACG |
| 1-GBA-R | CACTATGCAGAGGGGAAGTGGAAGG |
| 20-PARK2-F | TGCCCCATGCAGTTAATGTTTTTGA |
| 20-PARK2-R | CGGCTCACAAGCTTACAGAGAGTCG |
| 11-PARK2-1F | CAGCTCAGTATCTTGCATTTGCTGCTA |
| 11-PARK2-1R | TTGCTGTGTGCCGTCAGAGTAATAAAA |
| 11-PARK2-2F | CCATCTATTTCGTCTTTCCTGCCTCAT |
| 11-PARK2-2R | TTCAACATGAGTTTCCTTCCCCTTT |
| 21-PARK2-F | AGTGACTGGCTTCTATCTGGGTTCACA |
| 21-PARK2-R | ACTGTGGACTTTTCTCTGAGCATACCC |

**ddPCR**

| | |
|---|---|
| PARK2-E1-F | GAGGCGTGAGGAGAAACTAC |
| PARK2-E1-R | GGCTCTCCTGGGTTAAATCC |
| PARK2-E2-F | GAGGTCGATTCTGACACCAG |
| PARK2-E2-R | GTCCAGTCATTCCTCAGCTC |
| PARK2-E3-F | TCAGCAGAGCATTGTTCACA |
| PARK2-E3-4 | TCAGTGTGCAGAATGACAGC |
| PARK2-E4-F | CGGGAAAACTCAGGGTACAG |
| PARK2-E4-R | CATGCTGACACTGCATTTCC |
| PARK2-E5-F | CCATCTTGCTGGGATGATGT |
| PARK2-E5-R | TGACCAGGTACTTACTGCAC |
| PARK2-E6-F | AGTCGGAACATCACTTGCAT |
| PARK2-E6-R | TCCCCAGGAAAGAGAGTTCA |
| PARK2-E7-F | AGCCCCGTCCTGGTTTTC |
| PARK2-E7-R | GAGTAGCCAAGTTGAGGGTC |
| PARK2-E8-F | AGGATTCTGGGAGAAGAGCA |
| PARK2-E8-R | AGCATGGTTTTCTTCCCCAT |
| PARK2-E9-F | CAGTATGGTGCAGAGGAGTG |

**ddPCR (continued)**

| Primer | Sequence (5' to 3') |
|---|---|
| PARK2-E9-R | GTGACTTTCCTCTGGTCAGG |
| PARK2-E10-F | CCGGGAATGTAAAGAAGCGT |
| PARK2-E10-R | AGTTGTTCCTGAGGCTTCAA |
| PARK2-E11-F | AGGCCTACAGAGTCGATGAA |
| PARK2-E11-R | TTTCCACTGGTACATGGCAG |
| PARK2-E12-F | CCACTGGTTCGACGTGTAG |
| PARK2-E12-R | AGGTAGACACTGGGTATGCT |
| GBA-E1-F | GAGGCAAAACGAAATCCCAC |
| GBA-E1-R | GCATGAGTGACCGTCTCTTT |
| GBA-E2-F | GAAAACTCCATCCCCTCAGG |
| GBA-E2-R | CGGAATTACTTGCAGGGCTA |
| GBA-E3-F | TTGACTCACTCACCTGATGC |
| GBA-E3-R | AGTAGTTGAGGGGTGGAGAG |
| GBA-E4-F | CATACTCAGCTCCATCCGTC |
| GBA-E4-R | TCTGCAATGCCACATACTGT |
| GBA-E5-F | GTAGCAAATTTGGGCAGGG |
| GBA-E5-R | ATTCCCTGTGGATGTCCTCA |
| GBA-E6-F | ACAAGCAGACCTACCCTACA |
| GBA-E6-2-F** | ACAAGCAGACCTACCCTACC |
| GBA-E6-R | CTATGCAGACACCCCTGATG |
| GBA-E7-F | TCAGCATGGCTAAATGGGAG |
| GBA-E7-R | TTGGCTCAAGACCAATGGAG |
| GBA-E8-F | TTGTGGTGAGTACTGTTGGC |
| GBA-E8-R | TTACAGTTCTGGGCAGTGAC |
| GBA-E9-F | AAAGAGCATGGTGTTGGGGA |
| GBA-E9-R | CAGACCCAGAAGCAGCTAAA |
| GBA-E10-F | ACTGTCGACAAAGTTACGCA |
| GBA-E10-R | TCTCCCACATGTGACCCTTA |
| GBA-E11-F | GTCCAGGTCGTTCTTCTGAC |
| GBA-E11-R | TGGGTGGGTGACTTCTTAGA |
| GBA-E12-F | GGGAAAGTGAGTCACCCAAA |
| GBA-E12-R | GGCTTCCTGGAGACAATCTC |
| RPPH1-F* | AATGGGCGGAGGAGAGTAGTCTGAAT |
| RPPH1-R* | CGAAGTGAGTTCAATGGCTGAGGTG |
| TERT-F* | GCACACCTTTGGTCACTCCAAATTC |
| TERT-R* | CCACATAGGAATAGTCCATCCCCAGAT |

### *Digital Droplet PCR (ddPCR)*

The BioRad QX200™ AutoDG™ Droplet Digital™ PCR System was used to perform ddPCR with manufacturer protocols. In brief, *Hind*III-HF restriction enzyme (NEB, MA, USA) was used to digest genomic DNA prior to droplet generation. A 20µL reaction volume was employed for droplet generation, containing 10uL of 2x Q200 ddPCR EvaGreen Supermix, 15ng digested genomic DNA, and 0.2µM primers (See Supplemental Methods). Next, PCR was conducted in cycling conditions using Bio-Rad's C1000 Touch Thermal Cycler as follows: 5 minutes (mins) at 95ºC for enzyme activation, 40 cycles of 30 seconds (sec) at 95ºC and 1 min at 60ºC for denaturation and annealing/extension, 5 min at 4ºC and 5 min at 90ºC for signal stabilization. The droplet reading was then performed to obtain concentrations of positive droplets (number of positive droplets per uL of reaction) for each PCR reaction using Bio-Rad QuantaSoft TM Software.

**<u>Supplementary Figure 1</u>**: **Selected study pedigrees**. (A) *Subject 2*, *GBA* c. p.E365K (p.E326K) and *LRRK2* p.G2019S. (B) *Subject 13*, *GBA* p.L483P (p.L444P) and *LRRK2* p.G2019S. (C) *Subject 1*, *GBA* deletion exons 2-8.

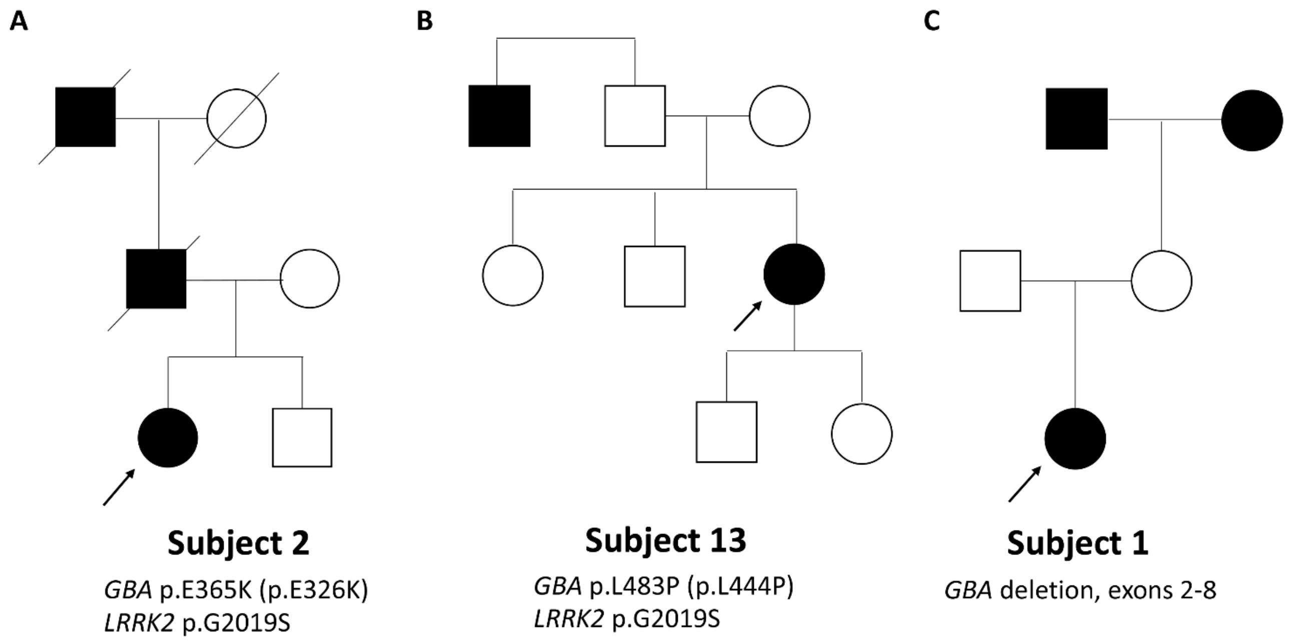

**<u>Supplementary Figure 2</u>**: ***PARK2* ddPCR screening**. ddPCR was used to perform an exon-by-exon screen for copy number variation involving one or more exons of *PARK2*. Primer pairs for the 12 exons of *PARK2* (*PRKN*) and two housekeeping genes *RPPH1* and *TERT* (i.e. neutral copy number in human genome) were used to obtain positive droplet concentrations from PCR in 92 individuals from the PD cohort for whom sufficient DNA was available. The y-axis shows the exon-by-exon results indicated by 13 columns with different colors, showing comparable ddPCR results to that of the average value of *RPPH1* and *TERT* used as controls. A y-axis value of 0.5 is consistent with a deletion, 1 indicates copy neutral (no deletion, no duplication), and 1.5 a duplication.

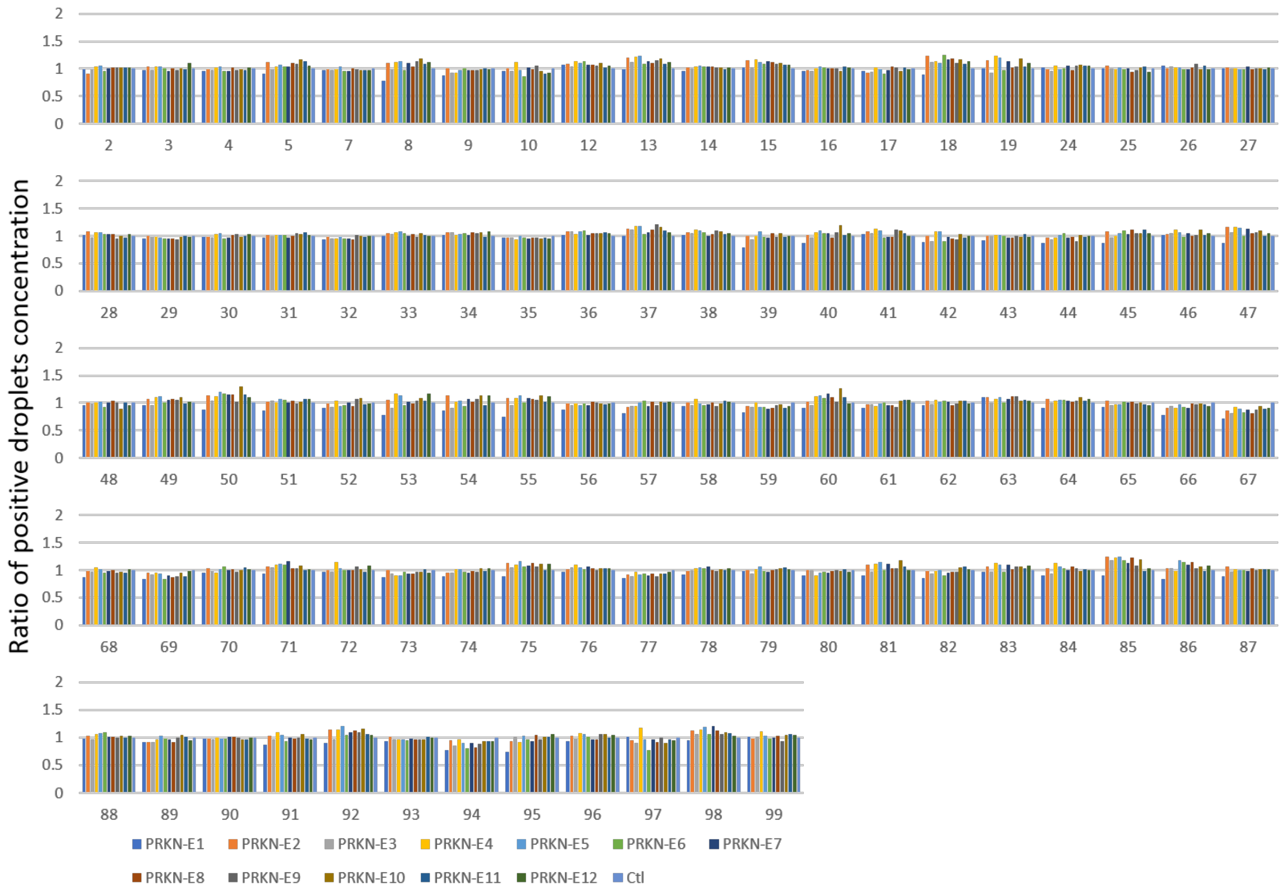

**<u>Supplementary Figure 3</u>**: ***GBA* ddPCR screening**. ddPCR was used to perform an exon-by-exon screen for copy number variation involving one or more exons of *GBA* in 80 individuals from the PD cohort for whom sufficient DNA was available. Primer pairs for 6 of the 12 exons of *GBA* produce amplicons concurrently from *GBA* and its pseudogene, *GBAP1*. This results in a doubling of the apparent number of exon copies identified by ddPCR for these exons: four, instead of two copies (ratio = 2) indicate copy number neutral for these exons.

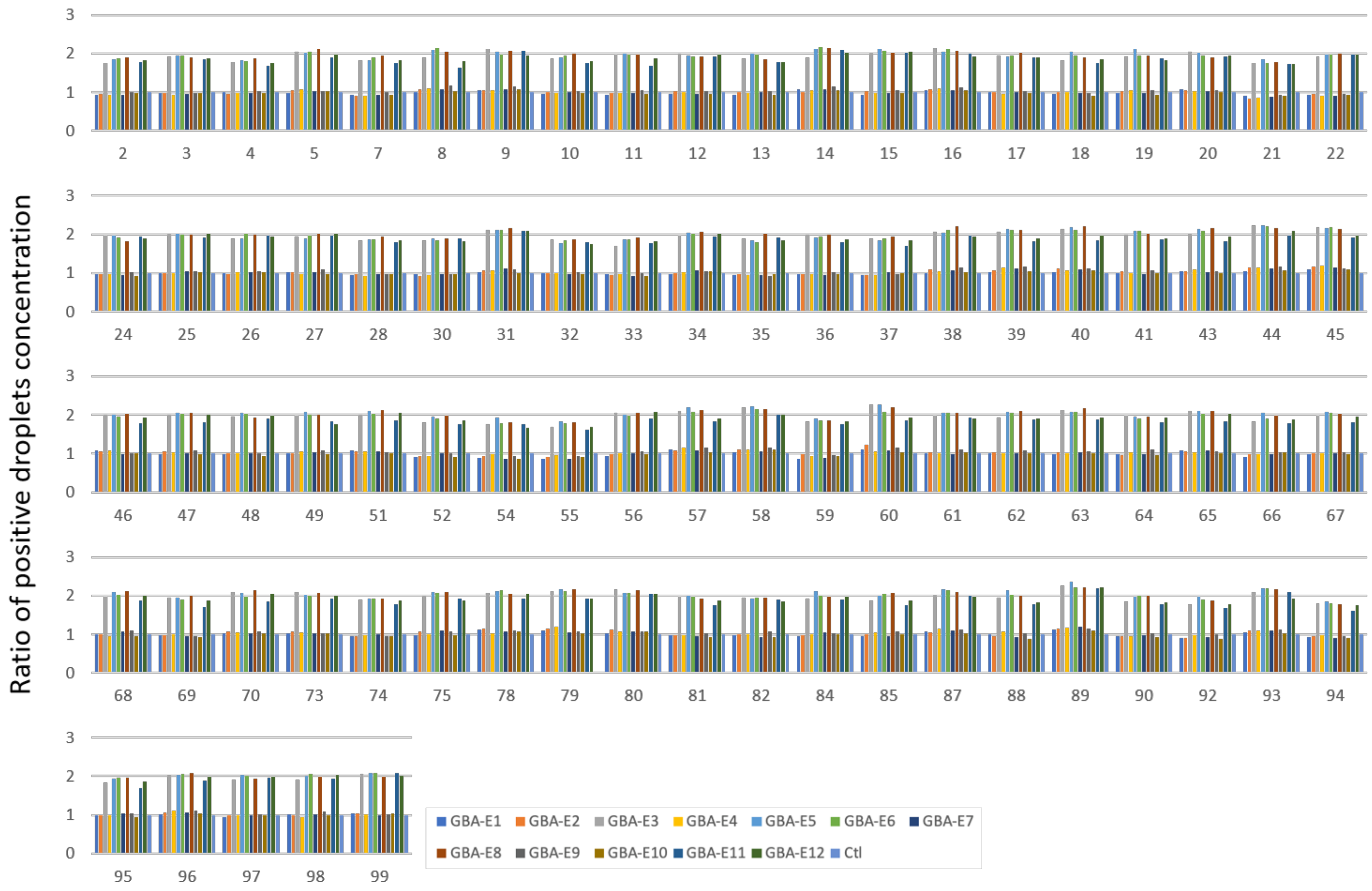

**<u>Supplementary Figure 4:</u> Amplification of exon 6 of *GBA* in subject 48.** A) Re-designed exon 6 primers incorporating the SNV chr1:155208250A>C (rs1317644130) into the primer sequence and Sanger confirmation. B) Subsequent ddPCR reactions using 1) initial primers on subject 48 (blue); 2) a combination of the initial and re-designed primers on subject 48 (orange); 3) initial primers on HapMap individual NA10851 shown as a control (gray).

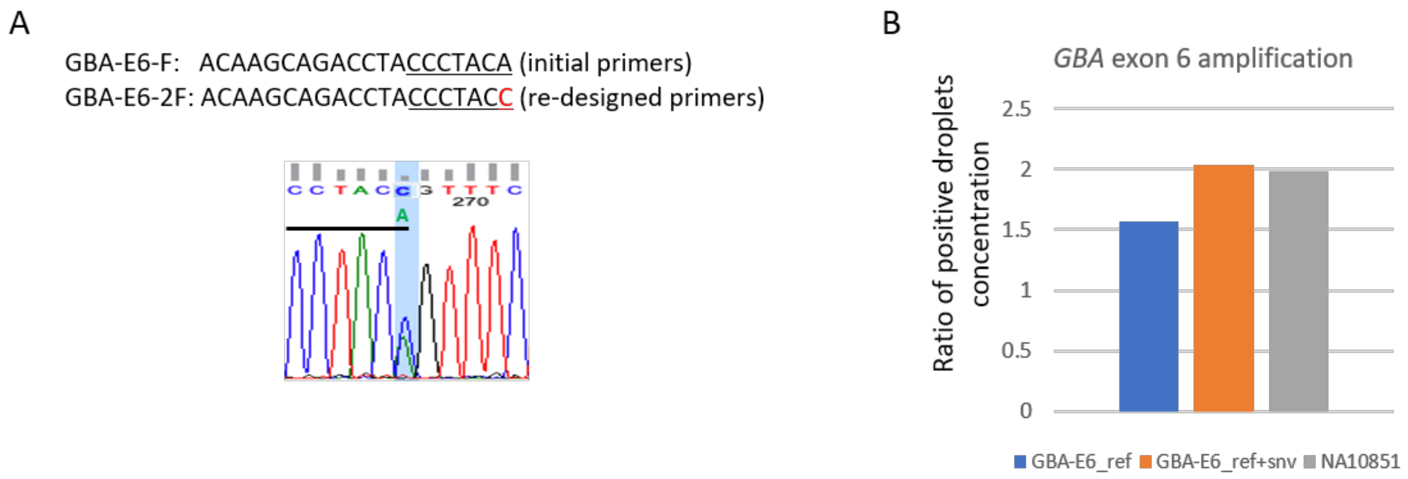


<u>Reference</u>:

Gu S, Posey JE, Yuan B, Carvalho CMB, Luk HM, Erikson K, et al. Mechanisms for the Generation of Two Quadruplications Associated with Split-Hand Malformation. Hum Mutat 2016; 37: 160–4.
